# Fitness compatibility and dengue virus inhibition in a Bangladeshi strain of *Aedes aegypti* infected with the *Wolbachia* strain *w*AlbB

**DOI:** 10.1101/2024.10.24.620043

**Authors:** Hasan Mohammad Al-Amin, Narayan Gyawali, Melissa Graham, Mohammad Shafiul Alam, Audrey Lenhart, Zhiyong Xi, Gordana Rašić, Nigel W. Beebe, Leon E. Hugo, Gregor J. Devine

## Abstract

Dengue cases in Bangladesh have surged in recent years. The existing insecticide-based control program, implemented in parts of the country, is challenged by issues of insufficient household coverage and high levels of insecticide resistance in the primary dengue virus (DENV) vector, Aedes aegypti. A more sustainable, effective alternative could be the implementation of a Wolbachia-mediated disease management strategy. Infecting mosquitoes with Wolbachia can change their reproductive compatibility and their ability to transmit DENV. These new phenotypes can be exploited to suppress or replace wild-type Ae. aegypti populations. Such strategies require the development of well-characterised Wolbachia-infected strains with biological characteristics that are comparable with local mosquitoes. We created and characterised a Wolbachia-infected Ae. aegypti strain with a Dhaka wild-type genetic background, and compared its reproductive compatibility, maternal inheritance, fitness, and virus-blocking ability to the parental strains (Dhaka wild-type and wAlbB2-F4). The new Ae. aegypti strain wAlbB2-Dhaka demonstrated complete cytoplasmic incompatibility with the wild-type strain and complete maternal transmission, retaining levels of pyrethroid resistance of the Dhaka wild-type (70% survival to 10 times the dose of permethrin expected to kill susceptible mosquitoes). No significant fitness costs were detected during laboratory comparisons of fecundity, fertility, survival, mating competitiveness, or desiccation tolerance. Compared to the wild-type strain, wAlbB2-Dhaka mosquitoes had a significantly reduced number of DENV genome copies in the bodies (44.4%, p = 0.0034); two-fold reduction in dissemination to legs and wings (47.6%, p < 0.0001); and >13-fold reduction of DENV in saliva expectorates (proxy of transmission potential) (92.7%, p < 0.0001) 14 days after ingesting dengue-infected blood. Our work indicates that the wAlbB2-Dhaka strain could be used for Ae. aegypti suppression or replacement strategies for dengue management in Bangladesh.

**Author summary:** Bangladesh is currently grappling with a series of severe dengue outbreaks, resulting in more than 300 thousand infections, 1700 fatalities, and a significant influx of patients requiring hospitalisation in 2023. These outbreaks coincide with the emergence of high levels of insecticide resistance in the Ae. aegypti population, which is contributing to the failure of conventional, insecticidal control strategies. In search of more sustainable, effective dengue control strategies, approaches using the endosymbiotic bacteria Wolbachia that render mosquitoes resistant to arbovirus infection are being trialled in a number of countries. We transinfected Dhaka wild-type Ae. aegypti with the Wolbachia strain wAlbB2 by backcrossing it with a Wolbachia wAlbB2-infected Ae. aegypti strain and assessed the new strain’s suitability for field release. We compared the fitness and virus-blocking ability of the backcrossed strain with its parental strains. We demonstrated that the new strain, wAlbB2-Dhaka, is as fit as the Dhaka wild-type strain and blocked ∼92% of dengue virus transmission. This strain has great potential for Wolbachia-mediated dengue management strategies in Bangladesh.

## Introduction

Dengue is the most common arboviral disease affecting humans, responsible for global epidemics and millions of infections [1]. The primary vector, *Aedes aegypti*, continues to proliferate globally, facilitated by globalisation, a heating climate, and urbanisation [2–5]. Dengue is a significant health concern for an estimated 3.9 billion people in 129 countries [6], with Asian countries carrying 70% of the dengue burden [1]. Bangladesh has recently experienced a series of severe dengue outbreaks, starting in 2019, with the worst outbreak to date occurring in 2023, with a total of 321,179 dengue cases and 1,705 deaths [7].

Global access to suitable vaccines or therapeutants for dengue control is still lacking. Managing the mosquito population remains critical for combating arbovirus transmission. Insecticidal applications are widely adopted but issues including limited household coverage, costs, and insecticide resistance undermine conventional insecticide-based *Ae. aegypti* control in Bangladesh [8]. Alternative insecticide classes might help mitigate the impacts of insecticide resistance, but this presents a suboptimal and short-term solution to the challenges of vector control [8]. Alternative dengue management tools are desperately required.

Over the past decade, the alphaproteobacteria *Wolbachia* has emerged as an important tool in controlling mosquito populations and the pathogens they transmit [9–13]. *Wolbachia* are obligate intracellular endosymbionts transmitted from mother to offspring during oogenesis, often with the highest density in ovarian germ cells. Their interaction with host insects ranges from parasitism to mutualism. *Wolbachia* transinfection into *Ae. aegypti,* which is not naturally infected with the bacterium, can be achieved through embryonic microinjection or through backcross mating with an infected colony [14–17]. Transinfection with certain *Wolbachia* strains can induce three highly exploitable phenotypes in the mosquito host: 1) maternal inheritance whereby infected female mosquitoes pass the bacteria to all of their offspring [18], 2) Cytoplasmic Incompatibility (CI) whereby crosses between *Wolbachia*-infected males and uninfected females, or females carrying a different strain of *Wolbachia*, results in embryonic death [16], and 3) virus-blocking, whereby *Wolbachia* infection can dramatically suppress virus infection, amplification and transmission [13]. These three traits allow *Wolbachia* to be used as a dengue control tool by either suppressing the native mosquito population through male-only releases [17, 19] or by replacing the local population with a virus-blocking phenotype through mixed releases of males and females [9, 20].

Release of *Wolbachia-*infected male and female *Ae. aegypti* as a population replacement strategy has been implemented in several countries, including Australia [9, 21], Indonesia [22–24], Vietnam [25], Brazil [11, 26], Colombia [27], and Malaysia [12, 20, 28]. *Wolbachia-* carrying male releases against *Ae. aegypti* (the incompatible insect technique, IIT), has been trialled in Australia [17], Singapore [29, 30], Mexico [31], and the USA [32]. Both replacement and suppression strategies require the released *Wolbachia-*infected mosquitoes to have equivalent fitness to local mosquitoes. In northern Queensland, Australia, a high prevalence of the *Wolbachia w*Mel strain (originally isolated from *Drosophila melanogaster*) has been maintained in the *Ae. aegypti* population for more than a decade since its release, and this has been associated with a 96% reduction in dengue incidence [9]. However, the prevalence of *w*Mel in *Ae. aegypti* after field releases has decreased in other countries [11, 33, 34]. One hypothesis for the loss of *Wolbachia* following releases is that the *w*Mel strain of *Wolbachia* can be affected by heat stress in hot climates [35]. The *w*AlbB *Wolbachia* strain (*Wolbachia* originating from *Ae. albopictus*) has been transinfected into *Ae. aegypti* [36–38] and forms a more robust infection under heat stress than the *w*Mel strain [39, 40]. Under simulated tropical environments (daily cycles from 27 °C to 37 °C), *w*AlbB infection remained at high densities in *Ae. aegypti* whereas *w*Mel was significantly reduced [41]. Under those conditions, the *w*AlbB infection was associated with greater inhibition of DENV infection and transmission [41].

In this study, we developed a *Wolbachia Ae. aegypti* Dhaka strain (*w*AlbB2-Dhaka) using a *Wolbachia* B-lineage strain (derived from *Ae. albopictus*) by backcrossing Dhaka *Ae. aegypti* wild-type male mosquitoes with high level of insecticide resistance with *w*AlbB2-infected *Ae. aegypti* females with an insecticide susceptible phenotype [17, 38]. We then characterised maternal inheritance, cytoplasmic incompatibility, virus-blocking, and biological fitness in comparison to the Dhaka wild-type.

## Results

### *Wolbachia* density in *w*AlbB2-Dhaka

*Wolbachia* was introduced into the Dhaka genetic background via a backcrossing procedure, where Dhaka wild-type males were mated with *w*AlbB-infected females. After six rounds of backcrosses, the mean density of *Wolbachia* infection in the final backcrossed colony, *w*AlbB2-Dhaka, was 6.01 log_10_ genome copies/µL (95% CI: 6 – 6.02 log_10_ copies/µL) and the mean density of *Wolbachia* in the parental *Wolbachia*-donor (*w*AlbB2-F4) colony was 6.04 log_10_ copies/µL (95% CI: 6.02 – 6.06 log_10_ copies/µL). There was no statistical difference in mean *Wolbachia* genome copies between strains (*T-test*, p = 0.07; t = 1.85; df = 94) (Fig 1A).

**Fig 1.**
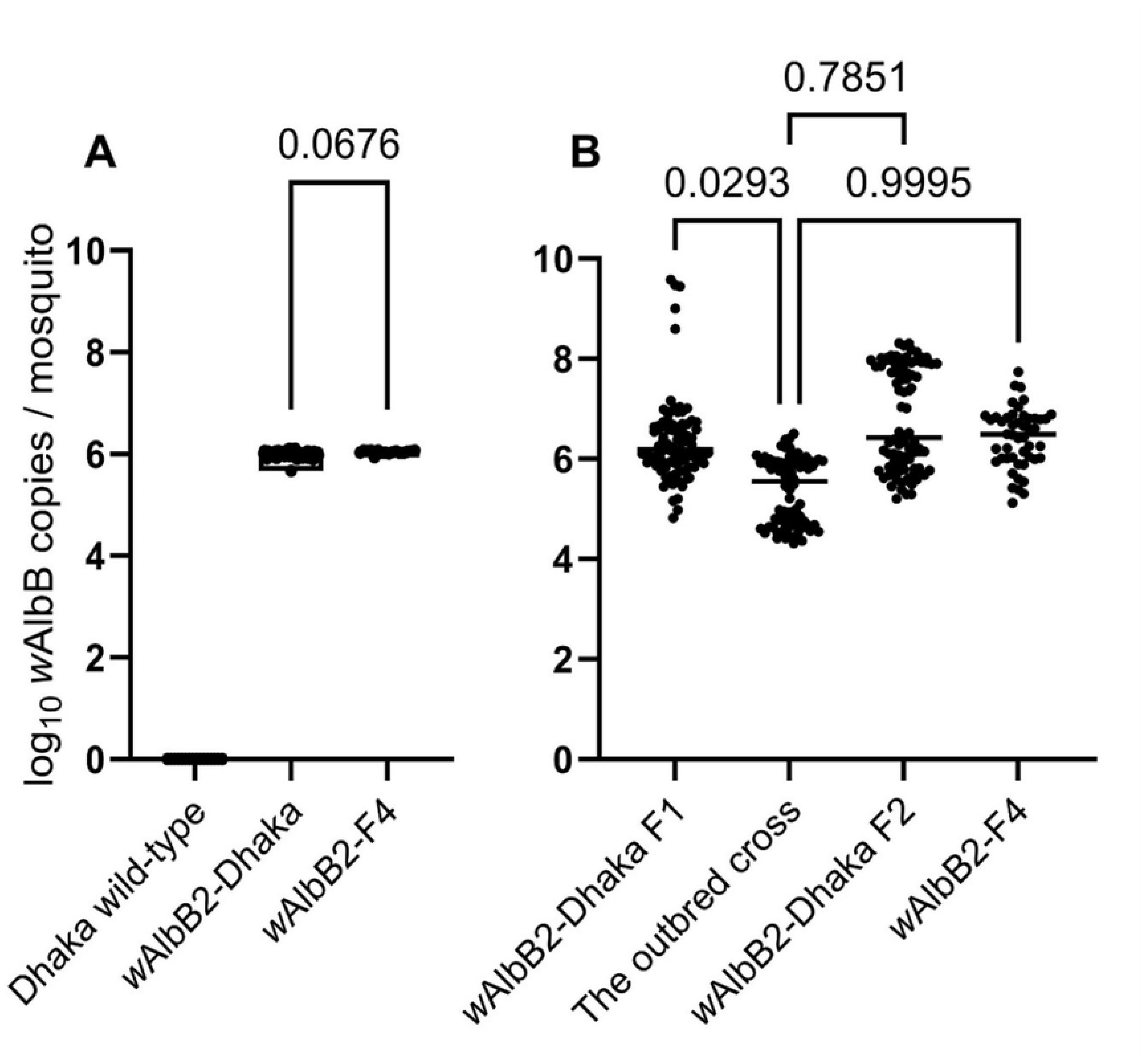
Genome copy numbers of *Wolbachia w*AlbB in parental and outcrossed Dhaka *Aedes aegypti* strains. (A) *w*AlbB copy number in *w*AlbB2-Dhaka and *w*AlbB2-F4 colonies. Comparative *w*AlbB concentration in *w*AlbB2-Dhaka (n = 80) and parental *w*AlbB2-F4 *Ae. aegypti* (n = 16). Dhaka wild-type was *Wolbachia*-free. (B) Maternal transmission of *w*AlbB by *w*AlbB2-Dhaka *Ae. aegypti*. All specimens of the outbred cross created by combining *w*AlbB2-Dhaka F1 and Dhaka wild-type mosquitoes were *w*AlbB positive demonstrating maternal transmission. The concentration of *w*AlbB in the outbred cross (n = 87) was compared with *w*AlbB2-Dhaka F1 (n = 80), *w*AlbB2-Dhaka F2 (n = 89), and parental *w*AlbB2-F4 (n = 48) by ANOVA with Dunnett’s multiple comparison tests.

### *w*AlbB2-Dhaka demonstrated complete maternal transmission

To test the maternal transmission, an outbred cross was created by combining *w*AlbB2-Dhaka F1 females (originated from *w*AlbB2-Dhaka sibling mating) and Dhaka wild-type males. The *w*AlbB infection density (genome copies) in the outbred cross was compared with *w*AlbB2-Dhaka F1, *w*AlbB2-Dhaka F2 (originated from *w*AlbB2-Dhaka F1 sibling mating), and the donor parental strain, *w*AlbB2-F4. All specimens were positive for *Wolbachia* (Fig 1B). The outbred cross had a lower *w*AlbB density than *w*AlbB2-Dhaka F1 (P = 0.03), however, no statistical difference was observed with *w*AlbB2-Dhaka F2 or *w*AlbB2-F4 (Fig 1B).

### Complete cytoplasmic incompatibility of *w*AlbB2-Dhaka strain

Complete (100%) CI was demonstrated when males from the fifth round of backcross (BC5), and from the final backcross colony (*w*AlbB2-Dhaka, after six rounds of backcrosses) were crossed separately with females of Dhaka wild-type. Crosses expected to have no CI, *i.e.,* crosses between males and females from the same strain (BC5, *w*AlbB2-Dhaka, Dhaka wild-type, and *w*AlbB2-F4 demonstrated 84 – 96 % larval hatch rates (S1 and S2 Figs.). Crosses between females of the Dhaka wild-type and males of the *Wolbachia* parental strain, *w*AlbB2-F4 (positive CI control), demonstrated complete CI (zero larval hatching, S1 and S2 Figs.).

### Phenotypic insecticide resistance in *w*AlbB2-Dhaka

We used CDC bottle bioassays to expose *w*AlbB2-Dhaka and Dhaka wild-type strains to 10 times the dose of permethrin that differentiates susceptible and resistant mosquitoes. High levels of resistance were demonstrated for both strains. Knockdown of *w*AlbB2-Dhaka and Dhaka wild-type was 41.75% (95% CI ± 6.63) and 36.75% (95% CI ± 4.86), respectively. No statistical difference was observed (T-test, p = 0.35; t = 0.97; df = 14) (Fig 2). Mortality of *w*AlbB2-Dhaka and Dhaka wild-type was 29.63% (95% CI ± 6.41) and 23.53% (95% CI ± 4.26), respectively. No statistical difference was observed (T-test, p = 0.08; t = 1.88; df = 14) (Fig 2). The *w*AlbB2-F4 colony was used as a susceptible control and tested with permethrin 1x. This colony exhibited complete susceptibility (Fig 2).

**Fig 2.**
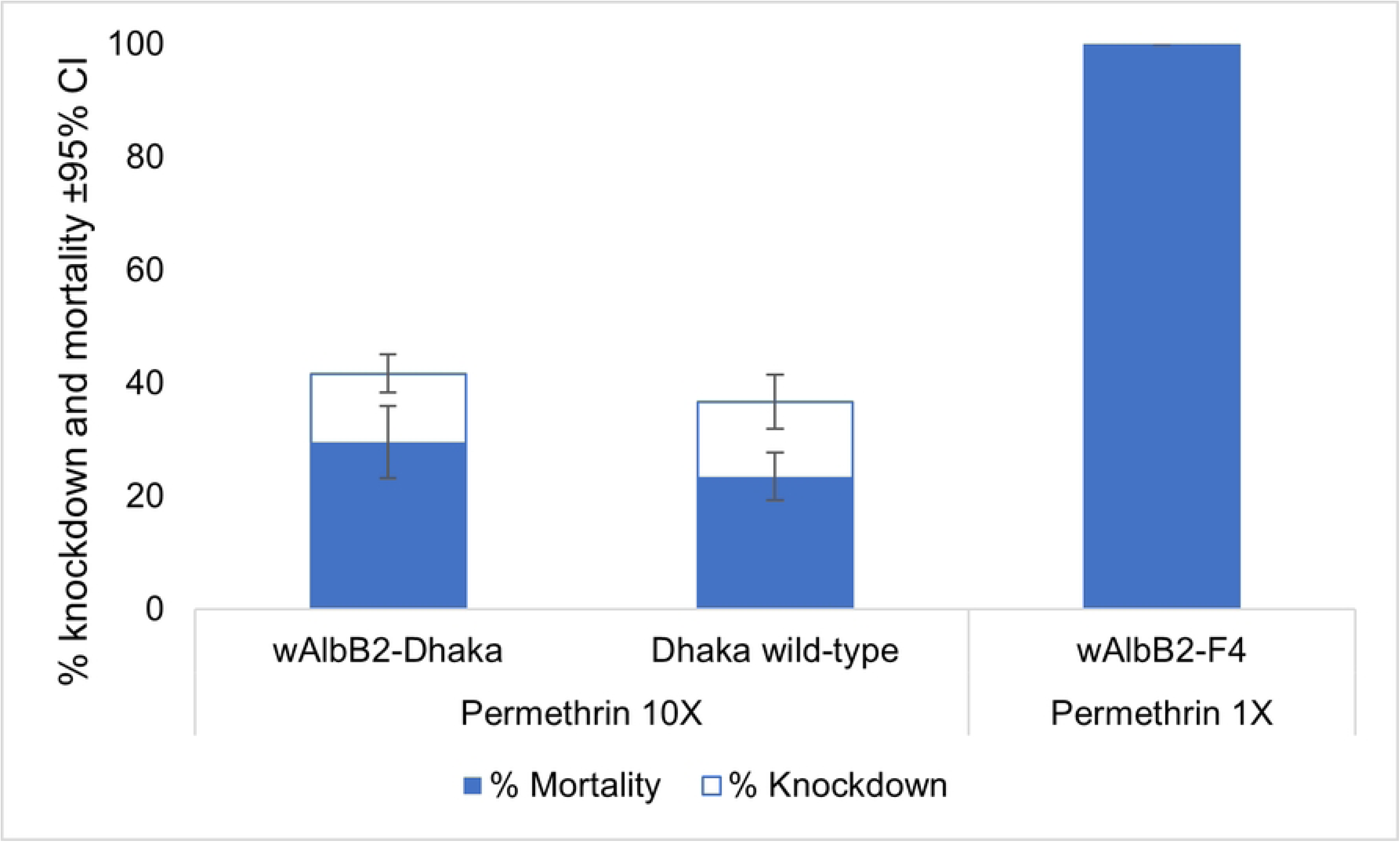
Compatible insecticide resistance phenotype maintained between the *w*AlbB2-Dhaka strain and Dhaka wild-type mosquitoes. Bioassay results (% knockdown and mortality ± 95% CI) of *w*AlbB2-Dhaka, Dhaka wild-type, and *w*AlbB2-F4 *Ae. aegypti*. Percent knockdown (white and blue combined) at 30 min and 24 h mortality (blue alone) is shown for mosquitoes exposed to 10x and 1x the diagnostic dose.

### Life history traits of *w*AlbB2-Dhaka female reproductive capacity

#### Fecundity, fertility and adult emergence

The mean number of eggs laid by *w*AlbB2-Dhaka females was 60.88 (95% CI: 26.84 – 64.91) and the mean for the Dhaka wild-type mosquitoes was 66.10 (95% CI: 57.17 – 75.03). There was no statistical difference between strains (*T-test*, p = 0.26; t = 1.14; df = 98) (Fig 3A). The mean percent hatch for *w*AlbB2-Dhaka eggs was 76.04% (95% CI: 73.53 – 78.56). For Dhaka wild-type eggs it was 78.67% (95% CI: 57.25 - 100). (*T-test*, p = 0.49; t = 0.69; df = 25) (Fig 3B). The mean percent emergence of *w*AlbB2-Dhaka and Dhaka wild-type was 91.46 (95% CI: 90.11 – 92.81) and 95.33% (95% CI: 89.08 – 101.60), respectively. (*T-test*, p = 0.06; t = 2.01; df = 25) (Fig 3C).

**Fig 3.**
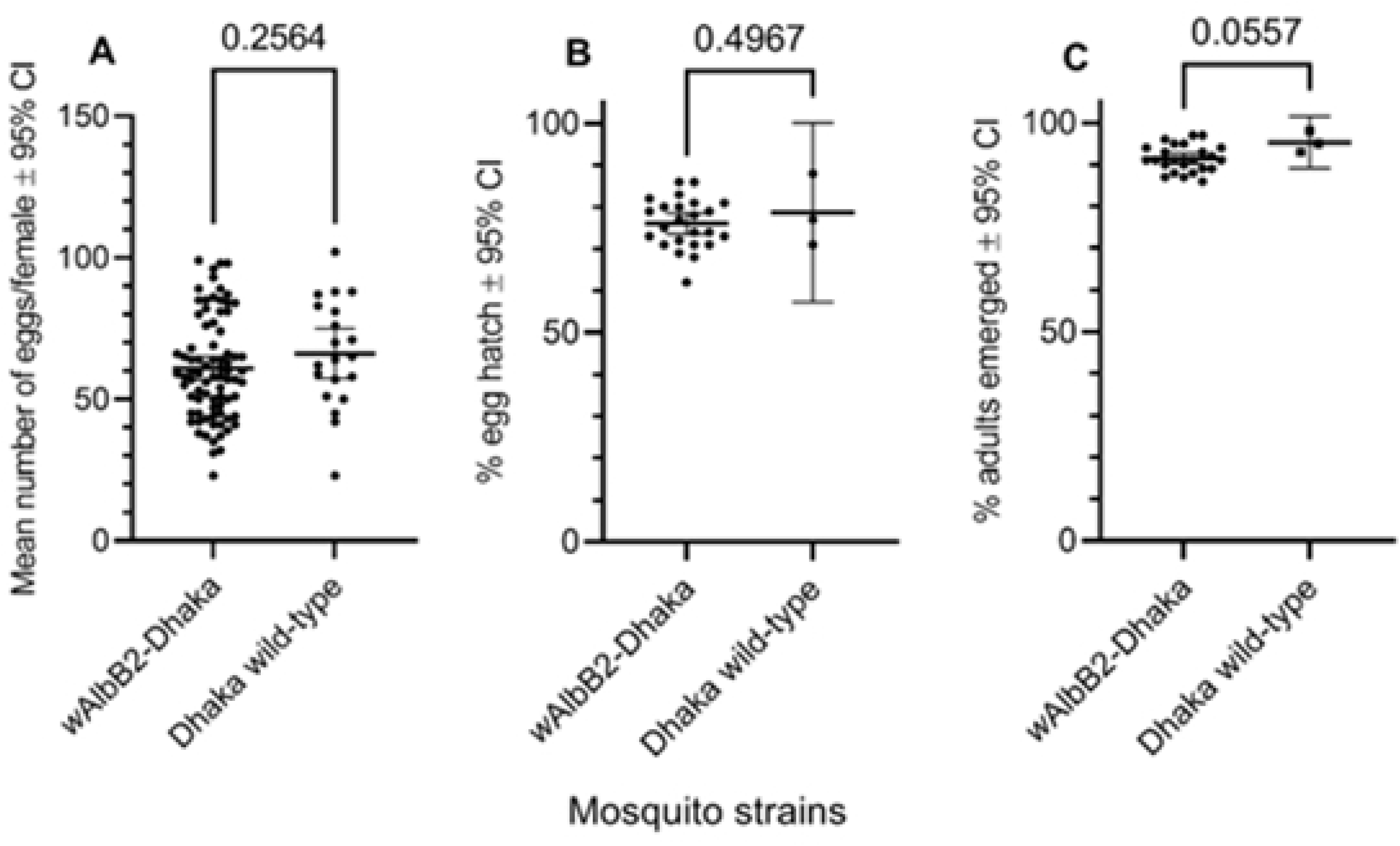
Life history traits of *w*AlbB2-Dhaka. (A) Comparison of female fecundity (number of eggs) between *w*AlbB2-Dhaka and Dhaka wild-type *Ae. aegypti*. (B) Comparison of female fertility (egg hatch rate) between *w*AlbB2-Dhaka and Dhaka wild-type *Ae. aegypti*. (C) Comparison of adult mosquito emergence from 100 4^th^ instar larvae between *w*AlbB2-Dhaka and Dhaka wild-type *Ae. aegypti* strains.

#### Survival of *w*AlbB2-Dhaka mosquitoes compared to the parental strains

The daily mortality of male and female mosquitoes of Dhaka wild-type, *w*AlbB2-Dhaka, and *w*AlbB2-F4 strains were compared. Survival of all female strains was similar over 28 days (Log-rank [Mantel-Cox] test; χ^2^ = 0.74; p = 0.69) (Fig 4A). Male survival was also comparable between strains (Log-rank [Mantel-Cox] test; χ^2^ = 3.48; p = 0.18) (Fig 4B).

**Fig 4.**
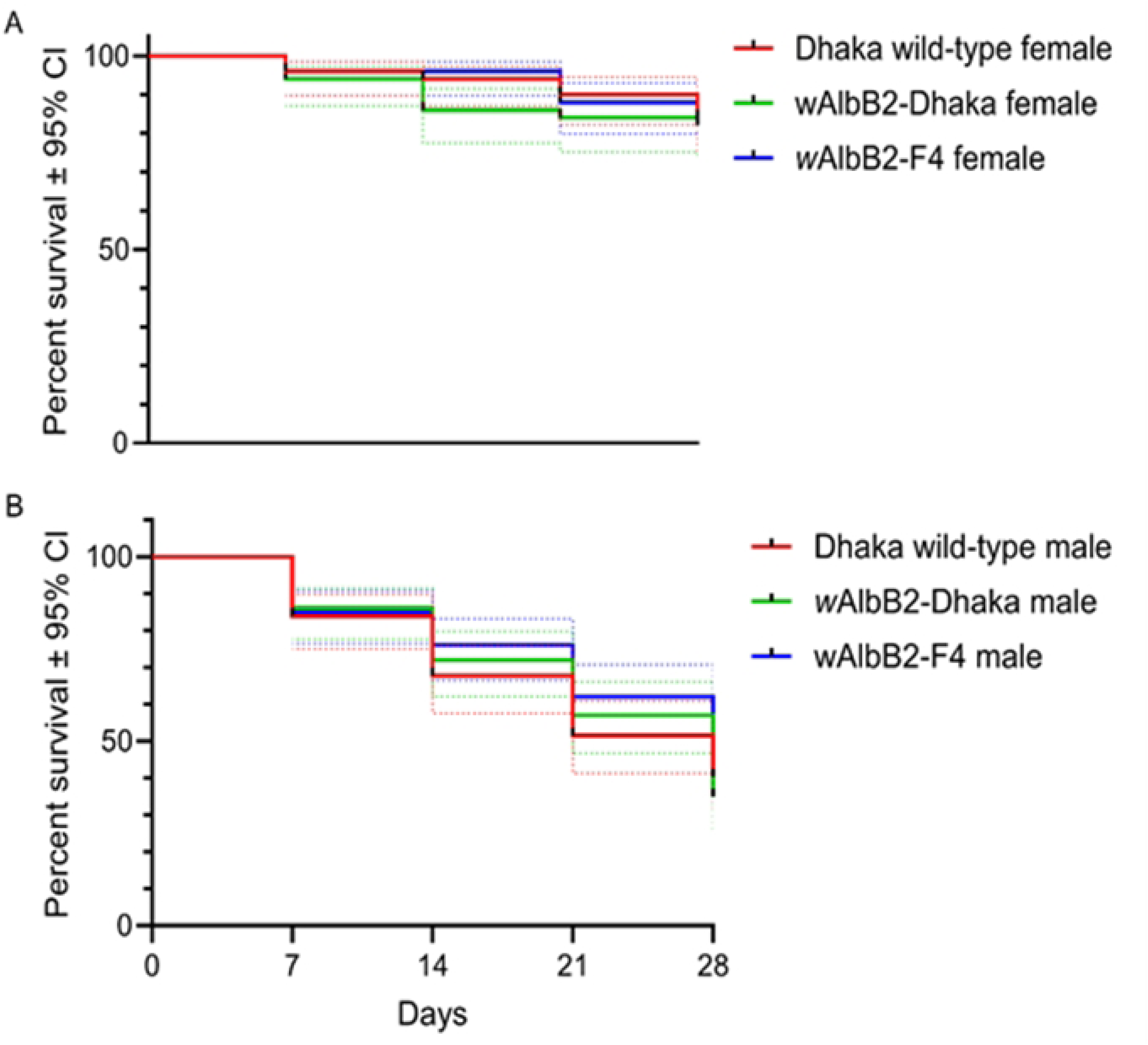
Equivalent life span of *w*AlbB2-Dhaka to the parental strains, Dhaka wild-type and *w*AlbB2-F4. Adult survival of Dhaka wild-type, *w*AlbB2-Dhaka, and *w*AlbB2-F4 *Ae. aegypti* females (A) and males (B). Survival analyses demonstrated no significant differences between the three strains for females or males. The dotted lines represent 95% CI.

#### Determination of egg quiescence (embryonic desiccation resistance)

The viability of dried, stored eggs was compared over time between *w*AlbB2-Dhaka F1 and parental Dhaka wild-type strains. Number of larvae and/or pupae was counted on day six of immersion of the stored eggs in water. No statistical difference (p > 0.05) was detected between strains for the weekly production of larvae/pupae (*T-test*; p = 0.96; t = 0.05; df = 30) (Fig 5). There was no difference in *w*AlbB density (log-transformed) in *w*AlbB2-Dhaka F1 and *w*AlbB2-Dhaka F2 females derived from eggs that had been stored for 4-16 weeks, respectively and neither diverged significantly from the *Wolbachia* density seen in freshly laid eggs of the donor strain *w*AlbB2-F4 (p > 0.05) (Fig 6).

**Fig 5.**
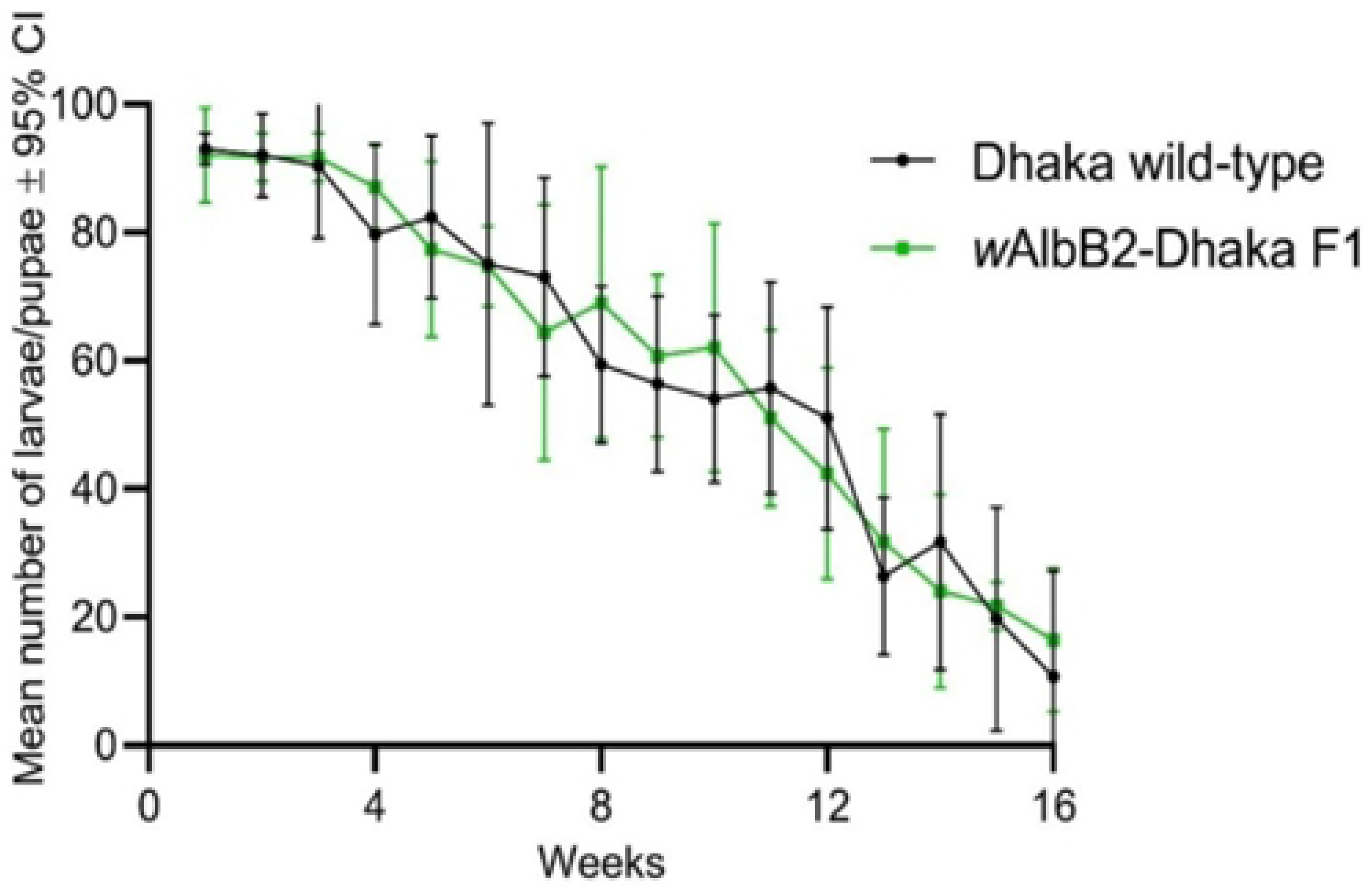
Egg viability of *w*AlbB2-Dhaka compared to the Dhaka wild-type *Aedes aegypti* strain. The number of larvae/pupae hatched from 100 eggs of Dhaka wild-type and *w*AlbB2-Dhaka F1 flooded weekly for up to 16 weeks were compared. Overall, no statistical difference was observed.

**Fig 6.**
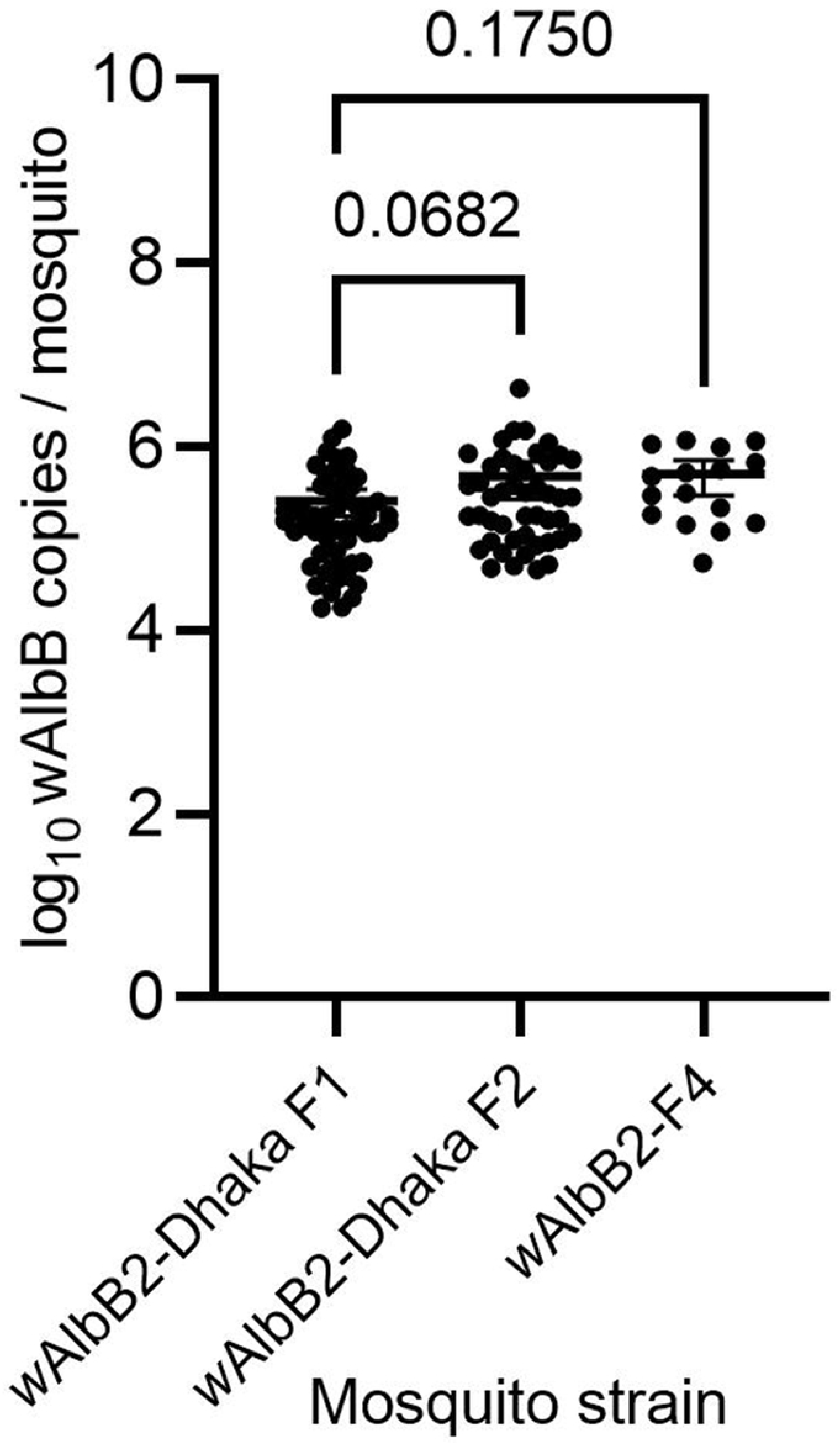
Effect of egg quiescence on *w*AlbB density in *w*AlbB2-Dhaka *Aedes aegypti*. *w*AlbB2-Dhaka F1 originated from 16-week-old eggs of *w*AlbB2-Dhaka F1, and *w*AlbB2-Dhaka F2 originated from 4-week old eggs from *w*AlbB2-Dhaka F2. Using ANOVA with Dunnett’s multiple comparison tests, no statistical differences were demonstrated in *w*AlbB density between *w*AlbB2-Dhaka F1 and *w*AlbB2-Dhaka F2 (p = 0.0682); or between *w*AlbB2-Dhaka F1 and parental *w*AlbB2-F4 (p = 0.1750).

#### Mating competitiveness

*Aedes aegypti* mating experiments were conducted in which male mosquitoes were ‘marked’ with the dye Rhodamine B () which renders sperm red before pairing the males and females. The presence of RhB-marked sperm in female spermathecae (sperm storage organs) indicated that the female first mated with an RhB-marked male mosquito. In one set of experiments, Dhaka wild-type female mosquitoes (n = 50) were allowed to mate with an equal number of RhB-marked *w*AlbB2-Dhaka males and unmarked Dhaka males. Following a period of mating, 48 Dhaka wild-type female mosquitoes were dissected. RhB can be transferred to female mosquitoes during mating events and therefore serves as a marker of mating. The spermathecae of 30/48 (62.5%) females contained RhB indicating successful mating with *w*AlbB2-Dhaka males (S3 A and B Fig.). Of the remaining females, the bursa was marked in one specimen, and 17 were unmarked (S3 C and D Fig.). These unmarked mosquitoes had, therefore, mated with Dhaka wild-type males as they contained sperm in the spermathecae (S1 Table). In a reciprocal experiment, Dhaka wild-type female mosquitoes (n = 50) were combined with an equal number of unmarked *w*AlbB2-Dhaka males and RhB-marked Dhaka males. A subset of Dhaka wild-type female mosquitoes (n = 34) were dissected and the spermathecae of 16/34 (47%) contained RhB and had therefore mated with male Dhaka wild-type mosquitoes. Of the remaining females, all 18 were unmarked and were considered to have mated with *w*AlbB2-Dhaka males (S1 Table). Fisher’s exact test showed no significant difference in the mating success of *w*AlbB2-Dhaka males (p = 0.18) between the two experiments.

#### Susceptibility of *w*AlbB2-Dhaka strain to dengue virus

To determine the pathogen blocking capability of *w*AlbB in Bangladeshi *Ae. aegypti*, the *w*AlbB2-Dhaka and parental strains were challenged DENV serotype 2 (DENV-2) at a titre of 1 × 10^6.9^ 50% cell culture infectious dose (CCID_50_) / ml via an infectious blood meal. Following a 14 d extrinsic incubation period, the presence and density of DENV-2 RNA in mosquito tissues was analysed using a qRT-PCR assay that detects a region of the DENV 3’ untranslated region (3’ UTR) [42]. Fourteen days after virus exposure, 96.4% (n = 28) of Dhaka wild-type mosquitoes had detectable DENV-2 RNA in bodies (carcass with wings and legs removed), whereas DENV was detected in only 53.6% (n = 28) of *w*AlbB2-Dhaka mosquitoes (44.4% reduction, Fisher’s Exact test, p = 0.0034) (Fig 7A). DENV-2 RNA was detected in 39.3% (n = 28) of *w*AlbB2-F4 mosquitoes, which was not significantly different from the *w*AlbB2-Dhaka strain (Fisher’s Exact test, p = 0.42) (Fig 7A). The median DENV-2 copy number in infected mosquitoes was 80% lower in *w*AlbB2-Dhaka compared to Dhaka wild-type (Kruskal-Wallis test, p < 0.0001). However, no significant difference was observed in DENV-2 copy numbers between *w*AlbB2-Dhaka and *w*AlbB2-F4 (Kruskal-Wallis test, p > 0.99) (Fig 7D).

**Fig 7.**
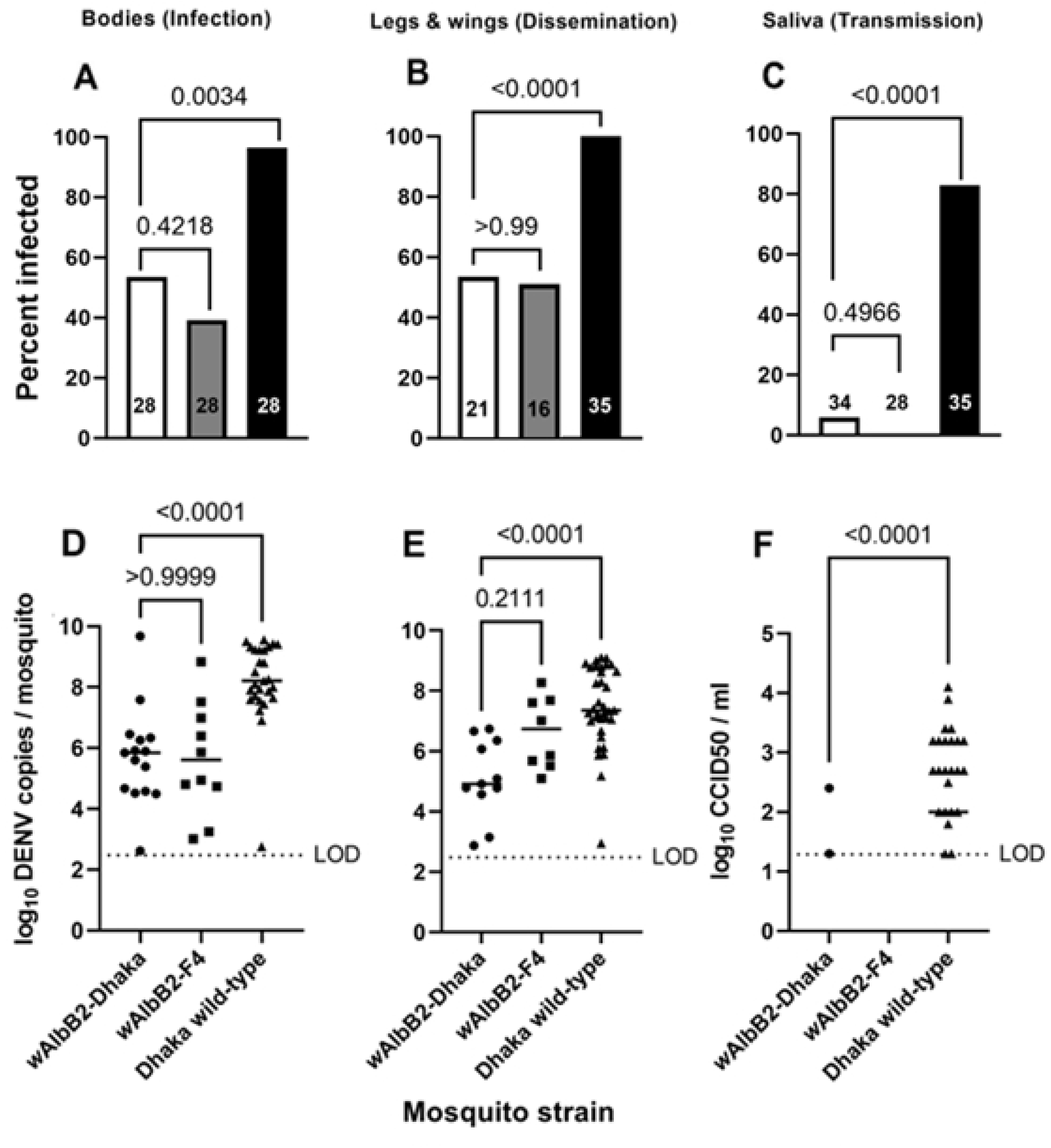
*Wolbachia* inhibits DENV-2 infections in *w*AlbB2-Dhaka mosquitoes. (A-C) DENV-2 virus prevalence in the bodies (A), legs and wings (B), and saliva (C) of *w*AlbB2-Dhaka, *w*AlbB2-F4, and Dhaka wild-type mosquitoes 14 days after feeding on a blood meal containing 1 × 10^6.9^ CCID_50_/ml (in C6/36 cells) of DENV-2 virus. P-values are shown for comparisons of mosquito infection rates between *w*AlbB2-Dhaka and Dhaka wild-type, and *w*AlbB2-Dhaka and *w*AlbB2-F4 (Fisher’s exact test). (D-F) DENV-2 infection intensity in DENV-2 positive mosquitoes in A-C. Virus copy numbers were determined from bodies (D), legs and wings (E) using qRT-PCR, and CCID_50_/ml was determined in saliva expectorates (F) using cell culture ELISA. P-values are shown for comparisons of median virus copy numbers between *w*AlbB2-Dhaka and Dhaka wild-type, and *w*AlbB2-Dhaka and *w*AlbB2-F4 (Kruskal-Wallis test). Solid lines indicate medians. LOD, limit of detection.

Dissemination of DENV-2 into mosquito legs and wings was noted in 100% (n = 35) of body-positive specimens of the Dhaka wild type. In contrast, there was a highly significant reduction in the percentage of *w*AlbB2-Dhaka mosquitoes with virus detected in legs and wings (52.4%, n = 21, Fisher’s Exact test, p < 0.0001) (Fig 7B). Dissemination in DENV-2 positive *w*AlbB2-F4 mosquitoes (50.0%, n = 16) was not significantly different from *w*AlbB2-Dhaka (Fisher’s Exact test, p > 0.99) (Fig 7B). Median DENV copy numbers in legs and wings varied significantly between *w*AlbB2-Dhaka and Dhaka wild-type (Kruskal-Wallis test, p < 0.0001). There was a 99.9% reduction in DENV copy numbers in the legs and wings of *w*AlbB2-Dhaka compared to the Dhaka wild-type. However, no significant difference in DENV copy numbers in legs and wings was demonstrated between *w*AlbB2-Dhaka and *w*AlbB2-F4 (Kruskal-Wallis test, p = 0.2111) (Fig 7E).

The presence and quantity of live virus in saliva, as a proxy of transmission potential, was analysed by cell culture ELISA [43]. Almost 83% (29/35) of the saliva expectorates of the Dhaka wild-type were positive for DENV-2, whereas only 5.9% of *w*AlbB2-Dhaka expectorates (2/34) were positive (Fig 7C). The 92.7% reduction was highly significant (Fisher’s exact test, p < 0.0001). Furthermore, no DENV-2 infection was detected in *w*AlbB2-F4 saliva samples (n = 28) (Fig 7C). DENV-2 copy number in positive saliva expectorates was also highly significantly lower in *w*AlbB2-Dhaka compared to Dhaka wild-type (99.1% reduction in median DENV-2 copy number; Kruskal-Wallis test, p < 0.0001) (Fig 7F).

#### *Wolbachia* distribution in *w*AlbB2-Dhaka and *w*AlbB2-F4 strains

The *wsp* antibody staining demonstrated widespread *w*AlbB infection throughout mosquitoes with a variation of staining density among tissue types (Fig 8).

**Fig 8.**
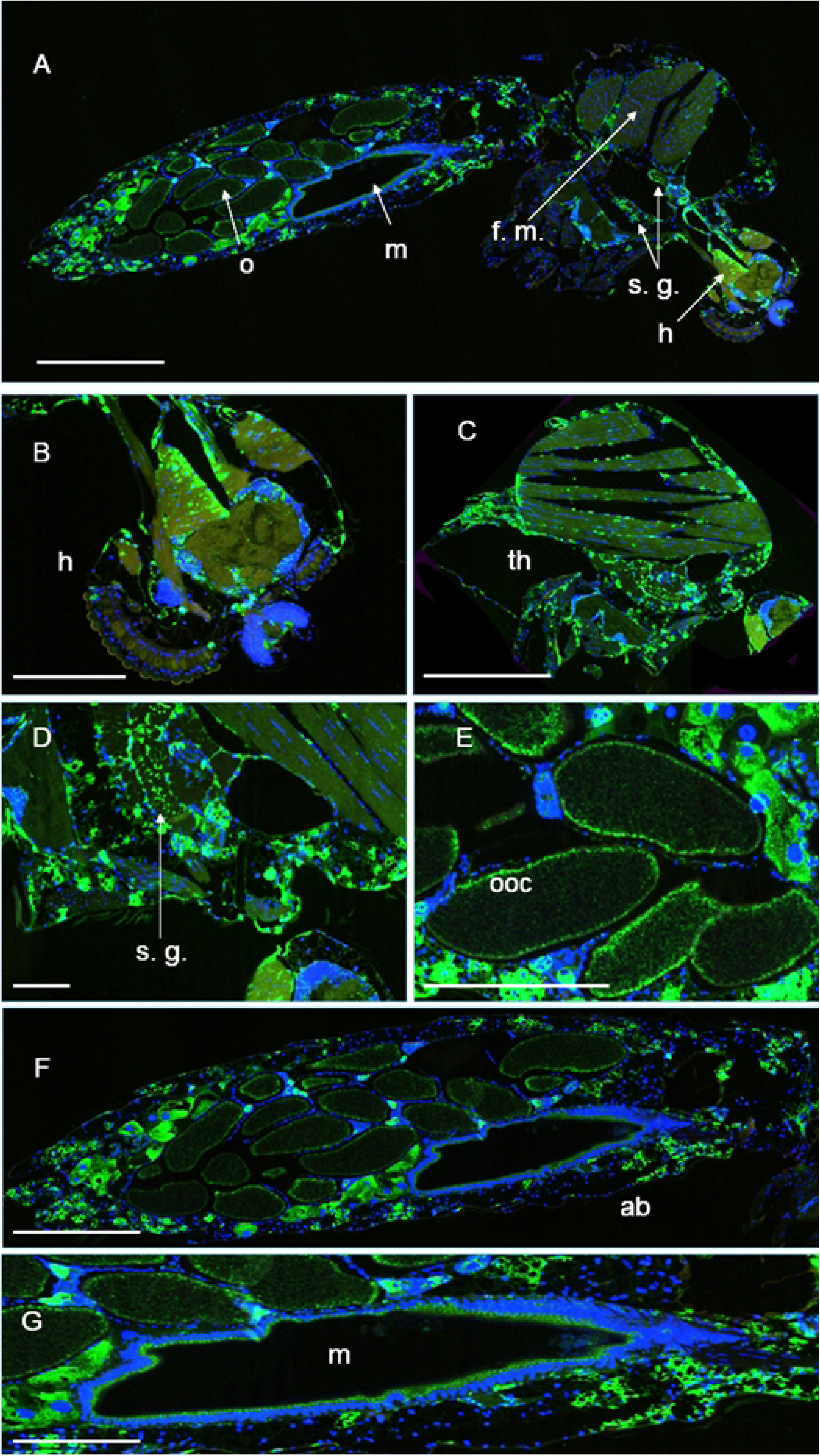
Histology of *w*AlbB infection in *w*AlbB2-Dhaka. Immunofluorescence analysis (IFA) using a rabbit polyclonal antibody against the *Wolbachia* surface protein (wsp) as the primary antibody and Alexa Fluor 488-conjugated donkey anti-rabbit antibody as the secondary antibody revealed the presence of *Wolbachia* infection across mosquito tissues. DNA was stained using DAPI. (A) Representative image of IFA staining in a midsagittal section through a whole mosquito. (B-G) High-resolution images displaying *Wolbachia* staining in the head, thorax, salivary gland, oocytes, abdomen, and midgut, respectively. Green: *Wolbachia*. Blue: DNA. h = head, s.g. = salivary glands, f.m. = flight muscle, m = midgut, o = ovary, ooc = oocytes, ab = abdomen. Scale bars: A and G 0.5mm, B-F 0.25mm.

The percentage of tissue samples positive for *Wolbachia* infection was compared between female *w*AlbB2-Dhaka and *w*AlbB2-F4 mosquitoes for different tissue types by Fisher’s exact test. The percentages of samples with *Wolbachia* infection did not differ between *w*AlbB2-Dhaka and *w*AlbB2-F4 strains for heads, thoraces, salivary glands, abdomens, and ovaries (S4 Fig).

#### DENV distribution within wild-type and *w*AlbB-infected strains

Dengue virus infection could be observed in tissues throughout the head, thorax, salivary gland, ovary, midgut, and abdomen of Dhaka wild-type mosquitoes following dual IFA, detecting both *Wolbachia* (green) and DENV (red) (Fig 9A-G). High DENV-2 staining densities were particularly observed in the midguts of Dhaka wild-type mosquitoes (Fig 9G). In contrast, for mosquitoes from the *Wolbachia-*infected strains (*w*AlbB2-Dhaka and *w*AlbB2-F4) that were infected with DENV-2, the virus was restricted to the midgut (representative example shown in Fig 9H).

**Fig 9.**
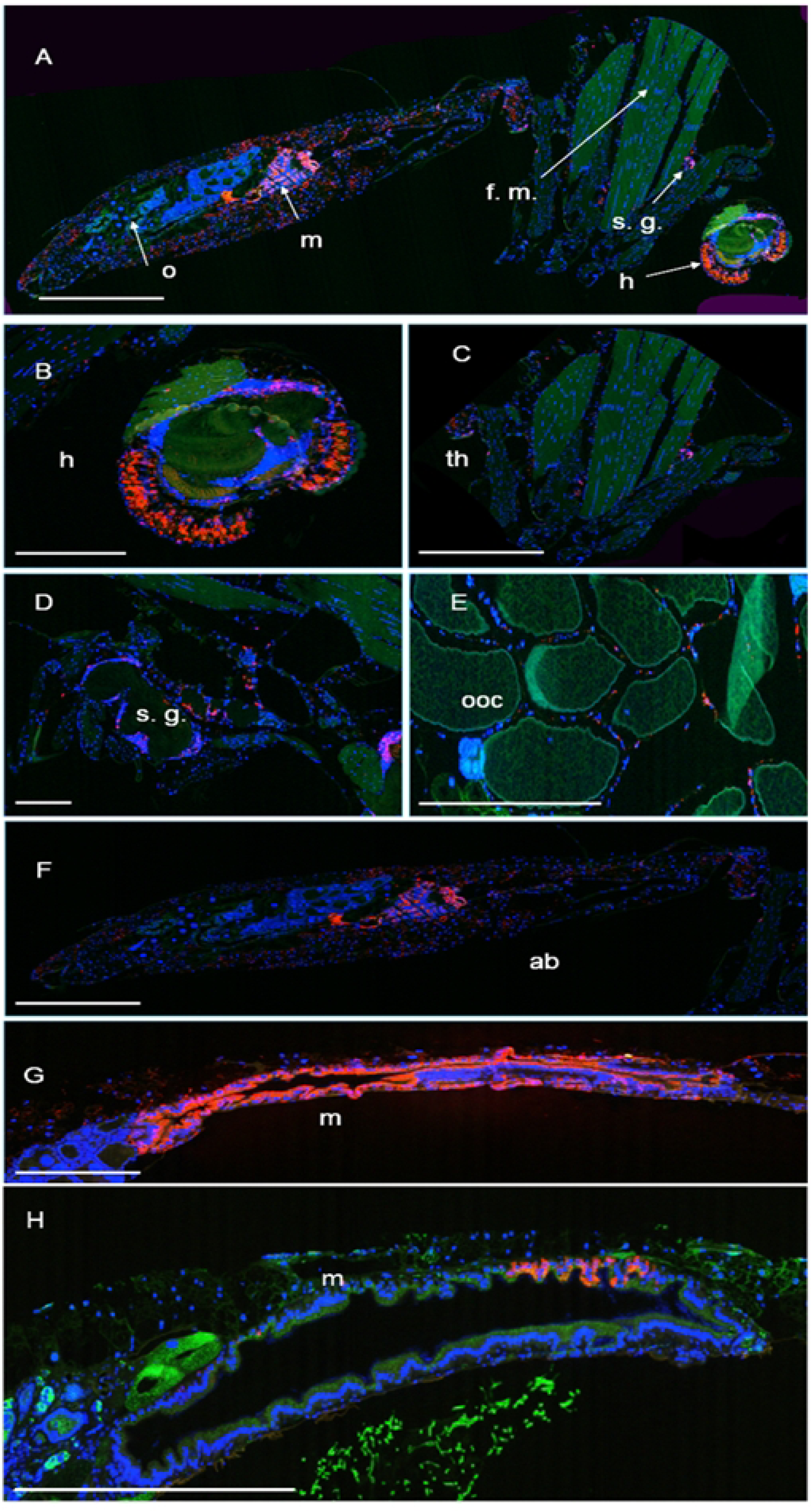
Dengue virus infection in Dhaka wild-type and *w*AlbB2-Dhaka. Fluorescent micrographs of mosquito midsagittal sections following dual immunofluorescence analysis of *Wolbachia* surface protein (wsp, green) and DENV-2 non-structural protein NS1 (red). (A-G) Example sections from Dhaka wild-type mosquitoes. (A) Whole-body section showing DENV-2 disseminated throughout the body. (B-G) High-resolution images of DENV-2 staining in the head, thorax, salivary gland, oocytes, abdomen, and midgut of Dhaka wild-type mosquitoes, respectively. (H) Example midgut of a *w*AlbB2-Dhaka mosquito infected with both *Wolbachia* and DENV-2. DENV-2 infection can be seen in midgut epithelial cells where *Wolbachia* was absent. Red: DENV-2. Green: *Wolbachia*. Blue: DNA. h = head, s.g. = salivary gland, f.m. = flight muscle, m = midgut, o = ovary, ooc = oocytes, ab = abdomen. Scale bars: A, G and H 0.5mm, B-F 0.25mm.

The proportion of different tissues infected with DENV was compared between *w*AlbB2-Dhaka, *w*AlbB2-F4, and Dhaka wild-type strains for different tissue types by IFA image analysis using Fisher’s exact test. Significantly higher DENV-2 infection prevalence was observed in Dhaka wild-type than *w*AlbB2-Dhaka or *w*AlbB2-F4 strains across all the tested tissues, including heads, thoraces, salivary glands, midguts, and abdomens (S5 Fig.).

#### *Wolbachia* density in *w*AlbB2-Dhaka and *w*AlbB2-F4 specimens used for vector competence study

The density of *Wolbachia w*AlbB was analysed in the specimens used for the vector competence study. The median *w*AlbB copy number did not differ significantly between *w*AlbB2-Dhaka and *w*AlbB2-F4 (Tukey’s multiple comparison tests, p = 0.97) in body tissues or in legs and wings (Tukey’s multiple comparison tests, p = 0.59) (S6 Fig.).

## Discussion

In this study, the *w*AlbB2-infected *Ae. aegypti* lab strain, *w*AlbB2-F4, was backcrossed with Dhaka wild-type *Ae. aegypti* to create a *w*AlbB2-infected *Ae. aegypti* strain, designated as *w*AlbB2-Dhaka, resulting in the first introduction of *Wolbachia* into a Bangladeshi *Ae. aegypti* strain. The overall distribution and densities of *Wolbachia* infection in *w*AlbB2-Dhaka were comparable to the *w*AlbB2-F4 parental strain. We demonstrated a 92.7% reduction in the competence of the *w*AlbB2-Dhaka strain for DENV-2 transmission compared to wild-type Dhaka *Ae. aegypti.* The backcrossed strain was also characterised in terms of insecticide resistance, CI, maternal transmission, and life history traits, including fecundity (egg number), fertility (egg hatch), adult emergence, daily survival, mating competitiveness, and egg viability. The *w*AlbB2-Dhaka *Ae. aegypti* demonstrated complete CI and maternal transmission, with equivalent fitness to Dhaka wild-type *Ae. aegypti* in caged laboratory comparisons, as well as equivalency with respect to permethrin resistance. Together with the known heat tolerance of the *Wolbachia w*AlbB2 strain [40], these represent ideal phenotypes for a *Wolbachia*-infected *Ae. aegypti* strain for deployment in Dhaka to counter the devastating outbreaks of dengue that have occurred in recent seasons.

Both the outbred cross (F1 progeny of female *w*AlbB2-Dhaka mated with Dhaka wild-type males) and the inbred cross (sibling mating between males and females of *w*AlbB2-Dhaka F1) demonstrated 100% maternal transmission, aligning with previous findings indicating complete maternal transmission of *w*AlbB in Australian, Indian, and Burkina Faso mosquito genetic backgrounds [44–46]. Recent laboratory experiments demonstrate the stability of maternal transmission of the *w*AlbB strains, even 15 years after the initial transinfection [47]. In addition, complete CI was demonstrated in our study when females were mated with Dhaka wild-type males after five rounds of backcrosses and when females from the final backcross colony *w*AlbB2-Dhaka were mated with Dhaka wild-type males. This generation of robust CI by *w*AlbB infection is consistent with findings from other studies [37, 45, 48, 49]. Cytoplasmic incompatibility serves as the cornerstone of *Wolbachia*-based mosquito control programs because: 1) it can be exploited in suppression campaigns where the mass release of *Wolbachia*-infected males can prevent the production of viable offspring [17], and 2) in replacement campaigns, it enables the propagation of *Wolbachia* through the female line, replacing the native populations with mosquitoes refractory to arbovirus infection [9, 12].

For *Wolbachia*-infected strains to survive and mate successfully with their wild-type targets, they must have comparable fitness to those local populations. Therefore, the insecticide-resistant phenotype of *w*AlbB2-Dhaka must be similar to the highly pyrethroid-resistant native Dhaka mosquitoes for the strain to be applied in local field deployments [8, 50]. Discrepancies in resistant phenotypes can prove problematic in *Wolbachia* release programs (e.g. in Rio de Janeiro, Brazil [51, 52]. We confirm that, after backcrossing, the novel *w*AlbB2-Dhaka strain demonstrated high intensity resistance to permethrin, reflecting the insecticide resistance profile of Dhaka wild-type *Ae. aegypti* populations.

Overall, no significant fitness cost of *w*AlbB infection in Dhaka *Ae. aegypti* was observed. *w*AlbB2-Dhaka showed comparable fecundity and fertility to the native Dhaka *Ae. aegypti*. In a recent study in India, *Wolbachia w*AlbB-infected females had significantly higher fecundity (number of eggs/female) than wild-type and *w*Mel-infected strains and there was similar fertility (egg hatch rate) among all strains [46]. Similar results were reported from Taiwan, where *w*AlbB infection in local *Ae. aegypti* caused no change in fertility [49]. One recent study indicated a significant loss of fecundity in *w*Mel-infected *Ae. aegypti* compared to the wild-type and the *w*AlbB-infected mosquitoes [53].

Mosquitoes released during *Wolbachia*-mediated control programs must demonstrate similar survival to local populations. For example, the prevalence of the *Wolbachia w*MelPop strain declined in *Ae. aegypti* populations following the cessation of deployments in Vietnam and Australia due to the induction of mosquito life-shortening by this *Wolbachia* strain [25]. Investigations of *w*AlbB have produced mixed results. *w*AlbB-infected mosquitoes have been reported to have reduced [44, 46] or equivalent [54] survival to wild-type mosquitoes. In our study, there was no significant difference in 28 d survival for males or females between Dhaka wild-type, *w*AlbB2-F4 or *w*AlbB2-Dhaka strains.

*Wolbachia* infection may affect egg viability over time [55, 56]. The *Wolbachia w*MelPop strain had a substantial negative impact on egg desiccation tolerance in an Australian *Ae. aegypti* strain, whereas *w*Mel infection in the same genetic host did not adversely affect desiccation tolerance [16, 44]. In the case of *w*AlbB, a 27-99% reduction in desiccation tolerance was previously recorded between 31-124 days, in comparison to an uninfected strain [44]. In our study, the viability of both Dhaka wild-type and *w*AlbB2-Dhaka F1 eggs were similarly reduced over 16 weeks of observation at 27 °C (± 0.5), 70% RH. At 10 weeks, the egg viability of *w*AlbB2-Dhaka and Dhaka wild-type remained > 50%. Although we did not compare *w*AlbB density throughout the 16-week storage period, comparison of *w*AlbB density in *w*AlbB2-Dhaka females reared from 4-week and 16-week-old eggs did not demonstrate any significant difference to fresh (one week old) eggs of *w*AlbB2-F4 under our laboratory conditions.

In earlier studies, a *w*AlbB-infected *Ae. aegypti* strain showed strong inhibition of DENV-2 in midguts and saliva expectorates [57] and both *w*Mel and *w*AlbB strains demonstrated similar reductions in DENV-2 viral loads [58]. In a study that characterised dengue blocking in our *Wolbachia* donor strain, *w*AlbB2-F4 [21], a four-fold reduction in DENV-2 genome copy numbers was detected in the bodies of *w*AlbB-infected females when compared with uninfected wild-type *Ae. aegypti*. In our study, we demonstrated a substantial reduction in DENV-2 dissemination to legs and wings and saliva. There was a 44.4% reduction in infection in bodies, a two-fold (47.6%) reduction in dissemination to legs and wings, and a >13-fold (92.7%) reduction in virus detections in *w*AlbB2-Dhaka saliva expectorate compared to the Dhaka wild-type strain. Dengue virus infection characteristics in the parental *Wolbachia*-infected *w*AlbB2-F4 were similar to the *w*AlbB2-Dhaka strain. A recent study from Taiwan demonstrated strong suppression of DENV-2 in *w*AlbB-infected backcrossed *Ae. aegypti* in the salivary gland and saliva samples at seven and 14-day post-blood meal compared to the wild-type strain [49]. In Malaysia, both lab and field-collected *w*AlbB-infected *Ae. aegypti* demonstrated significant reductions in DENV-2 infection in the salivary glands 12 days post-blood meal [59].

Male mating competitiveness is also essential for achieving effective invasion in population replacement and suppression strategies. *Wolbachia*-infected males must be as competitive as wild-type males in obtaining a mate. Our study demonstrated equivalent mating competitiveness of *w*AlbB-infected Dhaka male mosquitoes to Dhaka wild-type males in small cage experiments, which confirms results from other laboratory, semi-field and field studies [44, 49, 60].

In this study, double immunofluorescence staining of paraffin sections of *Wolbachia*-infected mosquitoes showed a dense distribution of *Wolbachia* throughout the body tissues including the head, thorax, salivary gland, abdomen, and ovary. The presence of *Wolbachia* in the developing eggs is typical of stable maternal transmission [61]. Fourteen days after mosquito exposure to the virus, the IFA studies detected few *w*AlbB2-Dhaka and *w*AlbB2-F4 individuals with DENV-2 in the gut cells that were not infected with *Wolbachia*. Dengue virus infection in *w*AlbB2-Dhaka and *w*AlbB2-F4 was mostly restricted to midguts, which had the lowest *w*AlbB density [38].

A limitation of this study was that reproductive traits and fitness determinations of the backcross strain *w*AlbB2-Dhaka were only tested under laboratory conditions due to strict biosecurity regulations. Experiments under semi-field and field conditions would provide more relevant data for predicting the results of field trials. No genetic analyses were undertaken to demonstrate similarity with the Dhaka parental strain. However, the backcrossing design followed was previously used to create the *w*AlbB2-F4 donor strain and resulted in >90% genetic similarity between wild-type parents after only four rounds of backcrosses [17, 62, 63]. We conducted six rounds of backcrossing, which can be expected to generate 98.44% genetic similarity.

In summary, this study showed that *w*AlbB infection in Bangladeshi *Ae. aegypti* reduced the vector competence by 92.7% and did not impose a detectable fitness cost. Our *w*AlbB2- Dhaka strain retained the high intensity pyrethroid resistance that is reflective of the local populations in Bangladesh. The virus inhibition and favourable reproductive fitness and life history traits provide encouraging results for potential pilot releases should this be amenable to regulators and the community. Dhaka is densely populated with high-density multi-storeyed apartments and hosts a prolific population of *Ae. aegypti* [64, 65]. A *w*AlbB-infected *Ae. aegypti* strain was shown to rapidly establish in multi-storeyed buildings in Malaysia and has been associated with a significant reduction in dengue outbreaks post-release [12, 20, 28]. Since current control tools are insufficient to control dengue in Bangladesh, the design of a *Wolbachia*-mediated strategy could address a major public health gap.

## Materials and methods

### Establishment of Dhaka *Ae. aegypti* colony

During June 2019, mosquito eggs were collected from five areas of Dhaka, Bangladesh using oviposition traps and shipped to QIMR Berghofer (QIMRB), Australia (import permit: 0002569896). Collections from one of the sites were not sufficient to establish a colony. These were harvested as fourth instar larvae (n = 40) and stored at -20 °C. F1 generations of the remaining four colonies were used for insecticide resistance phenotyping and genotyping studies. All colonies demonstrated high intensity resistance to pyrethroids (<98% mortality to 10 times the diagnostic dose) [8, 66, 67]. Eggs were then merged and reared to create the F2 generation and establish a single colony referred to as the Dhaka wild-type strain.

### Production of the *Wolbachia-*infected *Ae. aegypti* strain

The Dhaka F2 colony was backcrossed with the QIMRB laboratory-maintained *Wolbachia*-infected *Ae. aegypti* strain, *w*AlbB2-F4, to create the “*w*AlbB2-Dhaka” colony following the method of Yeap et al. [62] (S7 Fig.). In order to ensure virgin male and female mosquitoes for the backcross, pupae were manually sex-sorted based on size [68]. Male and female pupae were kept in separate cups with cotton wool soaked with 10% sucrose solution accessible to emerging adults through the mesh lids of the cups. Every 12 h, the sex of emerging adult mosquitoes was checked under a microscope on a chill table, and they were subsequently released into respective cups (male or female cups, n = 10 individuals/cup). If any individual of unexpected sex was identified from a cup, all adult individuals from the cup were discarded. When the desired number of male and female adults was collected in cups and maintained for 24 hours until the individuals were reproductively active [69]. Adult virgin mosquitoes (both male and female) were then released into rearing cages (1 ft^3^) (Bugdorm-1, MegaView Science Co. Ltd., Taichung City, Taiwan).

For each backcross, 200 Dhaka wild-type males were released with the same number of *w*AlbB2-F4 females. All mosquitoes were supplied 10% sucrose *ad libitum* and given a blood meal of defibrinated sheep blood (Serum Australis, NSW, Australia) supplied through a parafilm membrane after three days of confinement together [70]. Each cage of mosquitoes was provided with a moistened filter paper as a substrate for egg laying. Six generations of backcrossing were conducted to achieve the *w*AlbB2-Dhaka strain. All colonies were maintained at 27 °C (± 0.5), 70% RH, and 12:12 h day:night light cycling with 30 min dawn and dusk periods. By applying backcrossing for six generations, the *w*AlbB2-Dhaka is estimated to contain 98.44% genome similarity with Dhaka wild-type parents (S2 Table).

### Preparation of *wsp* standard for quantitative PCR

Quantitative PCR was conducted to assess *w*AlbB infection and concentration in the backcrossed individuals. Serial dilutions of a *Wolbachia* standard (*Wolbachia* surface protein gene [*wsp*]) were used to obtain a standard curve, and the absolute copy numbers for *Wolbachia* in a mosquito were obtained by comparing threshold cycle (CT) values with the standard curve. The standard curve was generated using a double-stranded plasmid containing a fragment of *wsp*. A set of forward and reverse cloning primers containing *Bgl II* and *Sal I* restriction endonuclease sites, respectively, were designed to amplify a 426 bp region of the *wsp* gene. The region was amplified by PCR cloned into plasmid pMDL2 using standard procedures. The supercoiled plasmid (designated as pMGL3_*wsp*) was transformed into DH10B competent *Escherichia coli* cells and purified using a Qiaprep spin miniprep kit (Qiagen). The plasmid was then linearised with *Sfi I*, purified using the Zymo DNA Clean and Concentrator kit (Zymo Research, CA, USA), and quantified using a NanoDrop spectrometer. Three readings were averaged, and copies/μl calculated. The plasmid was diluted to give 1 × 10^9^ copies/μl and serially diluted ten-fold down to 1 × 10^3^ copies/μl to generate a dilution series. Absolute quantification was conducted using a plasmid standard to determine the *Wolbachia* concentration within a mosquito [16] for 80 mosquitoes from each backcross generation. Mosquitoes from the parental strain *w*AlbB2-F4 were tested as positive controls [17, 38]. For the negative controls, *Wolbachia*-free colonies of *Ae. aegypti,* maintained at QIMRB (parental Dhaka wild-type) were used.

### Assessment of the maternal transmission of *w*AlbB2-Dhaka

The maternal transmission efficiency of the *w*AlbB infection was determined by testing 1-2 d old individual female offspring from crosses between *w*AlbB2-Dhaka F1 females and Dhaka wild-type males (the outbred cross) [47]. For each cross, one hundred 2-3 d old virgin *w*AlbB2-Dhaka F1 females were released into a cage with one hundred 2-3 d old virgin Dhaka wild-type males. A blood meal was offered and resulting eggs were air-dried and flooded after four days. To obtain *w*AlbB2-Dhaka F2, *w*AlbB2-Dhaka F1 mosquitoes were sibling-mated. Individuals from *w*AlbB2-Dhaka F1 (n = 87), emerged females from the outbred cross (n = 80), and *w*AlbB2-Dhaka F2 (n = 89) were tested for *w*AlbB infection and were compared with the *Wolbachia* infected strain *w*AlbB2-F4 (n = 48).

### Assessment of the CI of *w*AlbB2-Dhaka

Cytoplasmic incompatibility between *Wolbachia*-infected males and Dhaka wild-type females was characterised for backcross generation five (BC5; after five rounds of backcrossing) and the *w*AlbB2-Dhaka strain. Outcomes of mating (larval hatch) between BC5 males and Dhaka wild-type females and between *w*AlbB2-Dhaka males and Dhaka wild-type females were examined. Cytoplasmic incompatibility was assessed by observing the percentage of eggs that hatched successfully. Larval counts were made four days after egg flooding. Observations were compared to results from crosses where no CI was predicted (i.e., intra-strain crosses between male and female BC5, *w*AlbB2-Dhaka, *w*AlbB2-F4, and Dhaka wild-type). Positive controls included crosses between Dhaka wild-type females and *w*AlbB2-F4 males. During the experiments, 80 male BC5 and 80 female Dhaka wild-type pupae (1:1) were separated and kept in cups. The same was done with *w*AlbB2-Dhaka males and Dhaka wild-type females. For control sets, 10 male and female pupae (1:1) were separated. Sex sorting of pupae followed the methodology described above. After emergence, adults were sucrose fed (10%) and kept separate for 48 h, and then males and females were released together into a cage. After three days, 24 h sugar-starved females were offered defibrinated sheep blood (Serum Australis, Manila, NSW, Australia) as a blood meal, and engorged mosquitoes were allowed to oviposit for the next five days. Eggs were air-dried for four days under laboratory conditions [71], and one batch of 400 eggs for BC5 and *w*AlbB2-Dhaka each, and 50 eggs for each control set was flooded in reverse osmosis (RO) water with freshly ground fish food (TetraMin, Tetra GMBH, Melle, Germany) as a hatching stimulus [71]. Hatched larvae were counted on day four after flooding.

### Phenotypic characterisation of insecticide resistance in *w*AlbB2-Dhaka

A phenotypic characterisation of insecticide resistance was undertaken using standard bottle assays and exposing mosquitoes to 10 times the diagnostic dose of permethrin. The diagnostic dose (15 µg/bottle) should knock down all mosquitoes in a susceptible population at the diagnostic time of 30 min [66]. The knockdown phenotype (where mosquitoes are unable to stand or fly) was assessed 30 min post-exposure and mortality (if mosquitoes were unable to stand or were immobile) was assessed at 24 h [66, 67]. Each bioassay included ≥125 mosquitoes (≥25 mosquitoes per bottle with four bottles as biological replicates and one bottle as control). Bioassays were conducted using 10x the diagnostic dose of permethrin on *w*AlbB2-Dhaka and were compared with Dhaka wild-type and the susceptible *w*AlbB2-F4 *Ae. aegypti* strain (assayed at the diagnostic dose [1x] of permethrin). Control bottles were treated with the solvent acetone only.

### Determination of life history traits of *w*AlbB2-Dhaka female reproductive capacity

#### Number of eggs (fecundity)

Eighty blood-engorged, 5-8 d old mosquitoes from *w*AlbB2-Dhaka and twenty Dhaka wild-type were collected from the colony cages [49]. Mosquitoes were kept separately in clean plastic cups, and over the next five days, they were allowed to lay eggs on a wet filter paper substrate. After that, the filter papers were removed, and all eggs were counted under a microscope.

#### Egg hatch (fertility)

To assess female fertility (egg hatch) of *w*AlbB2-Dhaka in comparison to the Dhaka wild-type, 800 eggs (100 eggs in eight trays) of *w*AlbB2-Dhaka and 100 eggs of Dhaka wild-type colonies were flooded in separate trays. A larval count was conducted five days after egg flooding to assess the hatch rate. This experiment was replicated three times.

#### Adult emergence

To determine adult emergence rates, fourth instar larvae (n = 800; 100 larvae in eight trays) of *w*AlbB2-Dhaka, were observed until all had died or emerged as adults. The number of resulting adults was compared with Dhaka wild-type (n = 100). This experiment was replicated three times.

#### Survival

The lifespan of *w*AlbB2-Dhaka adults was compared with adults from the Dhaka wild-type and *w*AlbB2-F4 colonies. For this comparison, 100 male and female pupae were sex-sorted, and adult mosquitoes from each colony were kept in separate cages for four weeks (28 days). Mosquitoes were provided only 10% sucrose for sustenance and male and female mosquitoes were kept separately. Dead mosquitoes were counted weekly and removed from the cage [44].

#### Determination of egg quiescence

As the original bank of *w*AlbB2-Dhaka eggs had been exhausted by other experiments, *w*AlbB2-Dhaka F1 was used to compare with Dhaka wild-type in this experiment. Female mosquitoes of both strains were allowed to lay eggs on wet filter paper substrates for five days. Substrates were collected every 12 h to avoid any early hatching. These eggs were then stored in the insectary at 27 °C (± 0.5) and 70% RH. Three batches of 100 eggs from those *w*AlbB2-Dhaka F1 and Dhaka wild-type egg banks were flooded weekly for 16 weeks. Freshly ground fish food (TetraMin, Tetra GMBH, Melle, Germany) was added to the flooding trays as a hatching stimulus. On day six of flooding, larvae of all instars and any pupae that emerged were counted [44]. The presence of *Wolbachia* was also checked among the adult females (n = 54) that emerged from 16-week-old *w*AlbB2-Dhaka F1 eggs, and the *w*AlbB genome density was compared with females (n = 46) from 4-week-old *w*AlbB2-Dhaka F2 eggs, and females (n = 16) from freshly flooded eggs of parental *w*AlbB2-F4 colonies.

#### Mating competitiveness

To determine the mating competitiveness of *w*AlbB2-Dhaka males compared to Dhaka wild-type males, 50 *w*AlbB2-Dhaka and Dhaka wild-type males were fed with 0.2% rhodamine B [72, 73]. Rhodamine B is a thiol-reactive fluorescent dye that is used for tissue staining. The red-violet staining is visible under a fluorescence microscope. It is harmless to insects and has no impact on mating behaviour [74]. At the beginning of the experiment, pupae were sex-sorted, and virgin adults emerged into single-sex cages. Male mosquitoes were fed with 0.2% (w/v) RhB (Sigma Aldrich, 95% dye content, HPLC) dissolved in 10% sucrose (diluted in distilled water) for two days. This resulted in visible bright red staining, particularly on the abdomen [72]. Males were then introduced to the virgin female cages. All adult mosquitoes were 2-3 days old when they were combined for mating. Two sets of experiments were conducted. In the first, 50 RhB-marked *w*AlbB2-Dhaka males, 50 unmarked Dhaka wild-type females, and 50 unmarked Dhaka wild-type males were released into a cage (w30 x d30 x h30 cm) (S8 Fig.A). In the second, 50 unmarked *w*AlbB2-Dhaka males, 50 unmarked Dhaka wild-type females, and 50 RhB-marked Dhaka wild-type males were released in another cage (S8 Fig.B). For the positive control, 10 RhB-marked *w*AlbB2-Dhaka males were combined with 10 Dhaka wild-type females, and for the negative control, 10 unmarked *w*AlbB2-Dhaka males were combined with 10 Dhaka wild-type females.

Cages were left for 72 h with 10% sugar solution provided *ad libitum*. At this point, all females were considered mated [75], and were freeze-killed. The bursa and spermathecae of female mosquitoes were dissected into 1x PBS and then placed on a slide with a drop of fluorescent preservative containing DAPI (Fluoroshield with DAPI, Sigma Aldrich). The sample was gently crushed with a cover slip and placed under a fluorescent microscope (EVOS FL Auto, Life Technologies). Female mosquitoes receive sperm into their bursa, and after copulation, sperm travel to the spermathecae or are absorbed in the bursa [76]. If they receive a second insemination, sperm may stay in the bursa and are blocked from entering the spermatheca [75, 77]. In this experiment, a female mosquito was considered to be successfully mated by an RhB-marked male only if there was RhB in the spermathecae.

#### Dengue virus

The DENV serotype 2 (DENV-2) QML-16 strain was used in these experiments. This virus was initially isolated by Queensland Medical Laboratories (QML) from a dengue fever patient in Australia in 2015 [38, 78]. The virus was propagated in C6/36 cells at 30 °C, 5% CO_2_ for 5 days, and concentrated approximately 10-fold using an Amicon Ultra-15 centrifugal filter (Merck-Millipore, MA, USA).

#### Vector competence experiment Blood washing procedure

Defibrinated sheep blood was washed following the protocol of Reinhart et al. [79] (modified by Dr Narayan Gyawali, Queensland Health). Briefly, defibrinated sheep blood (Serum Australis, NSW, Australia) was added to a 50 ml centrifuge tube, and the volume was noted. The blood was centrifuged for 5 min at 2500 ×g. The plasma or supernatant was removed, and the blood was re-suspended in 5x volume of washing buffer (1% FBS [foetal bovine serum] in PBS at 37 °C). The supernatant was discarded, and the washing procedure was repeated three more times. Finally, the washed blood was re-suspended in re-constitution buffer (20% FBS in PBS at 37 °C) to the originally recorded volume.

#### Mosquito infection

A total of 60 female mosquitoes from each strain (*w*AlbB2-Dhaka, *w*AlbB2-F4, and Dhaka wild-type) were allocated into each of three 750 ml plastic containers with gauze lids. The mosquitoes were sugar-starved for 36 h prior to the experiment. DENV-2 virus stock was mixed with washed defibrinated sheep blood to a calculated titre of 1 × 10^7.3^ CCID_50_/ml. The blood virus mixture was offered to mosquitoes in glass artificial membrane feeders maintained at 37 °C for one hour [80]. A sub-sample of blood-virus mixtures was taken and stored before and after the feeding period to determine changes in titre. The mean DENV-2 titre fed was determined to be 1 × 10^6.9^ CCID_50_/ml in C6/36 cells. After feeding, mosquitoes were anaesthetised with CO_2_ and placed on a petri dish on ice. Fully engorged mosquitoes were returned to their respective containers and given access to a 10% sugar solution *ad libitum*. They were then housed in an environmental chamber (Aralab Fitoclima 600, aralab, Albarraque, Portugal) set to specific conditions, including a temperature of 28 °C, RH of 75%, and the same lighting conditions mentioned previously. Other non-fed and partially fed mosquitoes were discarded. At 14-d post-blood feeding, mosquitoes were anaesthetised and placed on ice. Mosquitoes were dissected to remove legs and wings, which were placed into 2 ml Eppendorf Safe-lock microcentrifuge tubes (Eppendorf, Hamburg, Germany) with three 2.3 mm zirconium silica glass beads. To collect saliva, mosquito bodies were placed on double-sided tape. A glass capillary tube containing 10 μl of saliva collection fluid, consisting of 10% FBS and 10% sugar, was then positioned over the proboscis of each mosquito for 20 min [81]. The contents of the mosquito saliva were expelled into a 1.5 ml microfuge tube. Finally, each mosquito body was transferred to a 2 ml safe-lock tube containing three beads [38].

#### Quantification of the virus from mosquito bodies and legs and wings

Virus nucleic acid was isolated using the Roche High Pure viral nucleic acid extraction kit (Roche Diagnostics GmbH, Manheim, Germany). The extraction process involved adding 200 μl of the working binding buffer (binding buffer mixed with poly[A] carrier RNA) to the tubes containing bodies or legs and wings, followed by tissue homogenisation through 1 min 30 s of shaking using a Mini Beadbeater-96 (BioSpec Products, Bartlesville, OK, USA). After centrifugation of the tubes at 8,000 × g for 1 min, 50 μl of Proteinase K was added, and the subsequent steps were carried out per the manufacturer’s protocol. One-step qRT-PCR was carried out using the TaqMan Fast Virus 1-Step Master Mix and primers and probe targeting the DENV 3’ untranslated region (UTR) region [42]. The reaction mix included 2.5 μl of 4 x Taqman Fast virus mastermix, 400 nM of each primer, 250 nM of the probe, and 1 μl of virus nucleic acid extraction in a total volume of 10 μl. The primers and probes used in the assay were synthesised by Macrogen (Macrogen, Seoul, Korea). Thermal cycling was carried out on a Corbett Rotorgene 6000 (QIAGEN/Corbett, Sydney, NSW, Australia), with an initial incubation at 50°C for 5 min, followed by denaturation at 95 °C for 20 s, then 40 cycles of 95 °C for 3 s, and 60°C for 30 s. Absolute quantification of virus copy number was performed using the Rotorgene 6000 software package, utilising a standard curve generated from 10-fold serial dilutions of a linearised plasmid containing the 3’UTR gene [42].

#### Cell culture enzyme-linked immunosorbent assay (ELISA) for determination of live virus in blood meals and mosquito saliva

Blood virus mixtures (pre-fed and post-fed) and saliva samples were titrated by cell culture enzyme-linked immunosorbent assay (ELISA) [82]. Blood meal samples were diluted by 10-fold serial dilution in virus media (RPMI 1640 supplemented with 2% foetal bovine serum [FBS, Thermo Fisher PTY, Australia], 0.1% ampicillin, and 1% penicillin-streptomycin [Gibco^TM^, Thermo Fisher, Australia]). For saliva samples, 50 µL virus media was added to saliva tubes (original 10 µL saliva expectorates), and the solution was mixed by pipetting. These were then centrifuged at 15,000 × g for 7 min at 4 °C. 57 µL of supernatant from each sample was then transferred to a sterile 1.5 tube, and centrifugation was repeated. 25 µL of supernatant was transferred to two wells in the first row of the 96-well ELISA plate (Corning Inc., Corning, New York, USA) containing 100 µL virus media per well. Five-fold serial dilutions were then performed. The contents of the dilution plate were transferred to equivalent wells of a second 96-well plate containing near-confluent C6/36 cell monolayers. Blood meal cell plates were incubated at 28 °C, 5% CO_2_, for five days, and saliva samples were incubated under the same conditions for seven days. The remaining procedure was conducted following Hugo et al. 2022 [38].

#### Histological analysis of *Wolbachia* and flavivirus infections

Immunofluorescence analysis (IFA) was used to detect *Wolbachia* and DENV-2 infections in thin paraffin sections of mosquitoes following established protocols [38]. Microscopy was performed using an Aperio ScanScope (Leica Biosystems) fluorescent microscope using filters for DAPI, Alexa 488 (FITC), and Alexa 555 (Cy 3), and exposure times of 0.125 s, 0.3 s, and 0.3 s, respectively [38, 43].

### Statistical analysis

Student’s *t*-test was used to compare *w*AlbB genome copies between *w*AlbB2-Dhaka and *w*AlbB2-F4. Dunnett’s multiple comparison tests of ANOVA were used to compare *w*AlbB concentration among *w*AlbB2-Dhaka F1, the outbred cross, and *w*AlbB2-Dhaka F2 with the parental *Wolbachia*-infected *w*AlbB2-F4. Insecticide resistance was classified according to the WHO guidelines where, for intensity assays at 10x the diagnostic dose, ≥98% mortality denotes moderate intensity resistance and <98% mortality indicates high intensity resistance (WHO, 2016). Student’s *t*-test was used to compare the percentage knockdown and mortality between *w*AlbB2-Dhaka and Dhaka wild-type and the proportion of *w*AlbB2-Dhaka with Dhaka wild-type for fecundity, fertility, and adult emergence. Survival curves for male and female mosquito strains were compared using a log-rank (Mantel-Cox) test. Comparisons between RhB-marked and unmarked *w*AlbB2-Dhaka male groups were made using Fisher’s exact test. Student’s *t*-test was also used to test the weekly and overall difference between *w*AlbB2-Dhaka and Dhaka wild-type mean egg viability. Dunnett’s multiple comparison tests of ANOVA were used to compare *w*AlbB concentration among *w*AlbB2-Dhaka F1, *w*AlbB2-Dhaka F2, and *w*AlbB2-F4. Dengue virus infection rates in body tissues between *w*AlbB2-Dhaka and Dhaka wild-type, and *w*AlbB2-Dhaka and *w*AlbB2-F4 were carried out using Fisher’s exact tests. Virus titres in body tissues of infected mosquitoes were compared between *w*AlbB2-Dhaka and Dhaka wild-type, and *w*AlbB2-Dhaka and *w*AlbB2-F4 using the Kruskal-Wallis test. For IFA analyses, the difference in *w*AlbB and DENV infection in body tissues between *w*AlbB2-Dhaka and *w*AlbB2-F4, and *w*AlbB2-Dhaka and Dhaka wild-type was compared using Fisher’s exact tests. The *w*AlbB copy number was compared between *w*AlbB2-Dhaka and *w*AlbB2-F4 using Tukey’s multiple comparison test. GraphPad Prism 9.4 (GraphPad Software, Boston, MA, USA) was used for these analyses.

## Supporting information

**S1 Fig. Cytoplasmic incompatibility after five rounds of backcross.** Results from the cross-mating experiments for CI of BC5 with Dhaka wild-type. Males of BC5 were combined with females of Dhaka wild-type. Eggs obtained from this colony were flooded (n = 400) and no larval hatch was observed (100% CI). No CI was observed in the negative controls, which consist of intra-strain crosses with BC5, Dhaka wild-type, and *w*AlbB2-F4. Positive control, i.e. a cross between males of *w*AlbB2-F4 and females of Dhaka wild-type, showed complete CI. BC = Backcross.

**S2 Fig. Cytoplasmic incompatibility of *w*AlbB2-Dhaka.** Results from the cross-mating experiments for CI of *w*AlbB2-Dhaka with Dhaka wild-type. Males of *w*AlbB2-Dhaka were combined with females of Dhaka wild-type. Eggs obtained from this colony were flooded (n = 400) and no larval hatch was observed (100% CI). No CI was observed in the negative controls, which consist of intra-strain crosses with *w*AlbB2-Dhaka, Dhaka wild-type, and *w*AlbB2-F4. Positive control, i.e. a cross between males of *w*AlbB2-F4 and females of Dhaka wild-type, showed complete CI.

**S3 Fig. Mating competitiveness of *w*AlbB2-Dhaka males.** Presence or absence of RhB on female spermathecae. (A) spermathecae of mated Dhaka wild-type *Ae. aegypti* female using bright-field microscopy. (B) the presence of RhB in the spermathecae shown in (A). (C) shows the spermathecae of an individual which has not been mated with a marked male, and (D) shows the absence of RhB. s, spermatheca. Scale bars = 0.2 mm.

**S4 Fig. Prevalence of *w*AlbB infection in different mosquito tissues.** Comparison of the proportion of tissue samples infected with *Wolbachia w*AlbB between *w*AlbB2-Dhaka, *w*AlbB2-F4, and Dhaka wild-type strains for different tissue types from the histological analysis. No *w*AlbB-infection was observed in Dhaka wild-type mosquitoes, and percentages of infected samples were not significantly different between *w*AlbB2-Dhaka and *w*AlbB2-F4 strains for any tissue (Fisher’s exact test).

**S5 Fig. Prevalence of DENV-2 infection in mosquito tissues.** Comparison of DENV-2 infection from the histological analysis between *w*AlbB2-Dhaka, *w*AlbB2-F4, and Dhaka wild-type. *w*AlbB2-Dhaka and *w*AlbB2-F4 demonstrated no significant differences (Fisher’s exact test, p > 0.05), whereas DENV-2 infection was significantly higher in Dhaka wild-type mosquitoes (Fisher’s exact test, p < 0.0001).

**S6 Fig. Density of *w*AlbB infection in mosquitoes.** *w*AlbB infection intensity in mosquitoes used in vector competence tests. *w*AlbB copy numbers were determined from bodies (n = 36 in all strains) and legs & wings (n = 21 in *w*AlbB2-Dhaka, n = 17 in wAlbB2-F4, and n = 35 in Dhaka wild-type) using quantitative PCR (qPCR). P-values are shown for comparisons of median *w*AlbB copy numbers between *w*AlbB2-Dhaka and *w*AlbB2-F4 (Tukey’s multiple comparison tests). Dhaka wild-type is a naturally *Wolbachia*-free mosquito strain. Lines indicate medians. LOD, limit of detection.

**S7 Fig. Backcross scheme to create *w*AlbB2-Dhaka *Ae. aegypti* strain.** Backcrossing was conducted between Dhaka wild-type males and *w*AlbB2-F4 females to establish the *w*AlbB2-Dhaka *Ae. aegypti* strain. In each backcross, the male and female ratio was 1:1 (n = 200 individuals/sex). BC = Backcross.

**S8 Fig. Mating competitiveness experiment.** Schematic diagram of the two combinations of marked and unmarked males used in mating competitiveness experiment. A: 50 RhB-marked *w*AlbB2-Dhaka males, 50 unmarked Dhaka F8 females, and 50 unmarked Dhaka F8 males. B: 50 unmarked *w*AlbB2-Dhaka males, 50 unmarked Dhaka F8 females, and 50 RhB-marked Dhaka F8 males.

**S1 Table. Mating competitiveness of *w*AlbB2-Dhaka males.** Comparative mating competitiveness between *w*AlbB2-Dhaka males and Dhaka wild-type males. No statistical difference in mating success was observed between males of *w*AlbB2-Dhaka and Dhaka wild-type strains.

**S2 Table. Calculation of Dhaka wild-type genome in *w*AlbB2-Dhaka.** Analysis of Dhaka wild-type male genome percentage in *w*AlbB2-Dhaka *Ae. aegypti* after six rounds of backcrosses.

## Acknowledgments

The *w*AlbB2-Dhaka strain was created under an MTA with the International Centre for Diarrhoeal Disease Research, Bangladesh (icddr,b). We express gratitude to the Histotechnology and Microscopy Facility in QIMR Berghofer for help in preparing slides and for providing training on fluorescent microscopy, respectively. We are also grateful to Stacey Llewellyn, QIMR Berghofer for assistance with statistical analysis.

## Author Contributions

**Conceptualisation:** Hasan Mohammad Al-Amin, Leon E. Hugo, Gordana Rašić, Nigel W. Beebe, Gregor J. Devine

**Data curation:** Hasan Mohammad Al-Amin

**Formal analysis:** Hasan Mohammad Al-Amin, Leon E. Hugo

**Funding:** This work was done in partial fulfilment of a PhD, made possible through a QIMR Berghofer International PhD Scholarship, the resources of the Mosquito Control Laboratory, QIMR Berghofer, and a tuition fee waiver from The University of Queensland.

**Investigation:** Hasan Mohammad Al-Amin

**Methodology:** Hasan Mohammad Al-Amin, Leon E. Hugo, Gregor J. Devine

**Resources:** Hasan Mohammad Al-Amin, Narayan Gyawali, Melissa Graham, Mohammad Shafiul Alam, Zhiyong Xi, Leon E. Hugo, Gregor J. Devine

**Supervision:** Leon E. Hugo, Gordana Rašić, Nigel W. Beebe, Gregor J. Devine

**Visualisation:** Hasan Mohammad Al-Amin

**Writing-original draft:** Hasan Mohammad Al-Amin

**Writing-review and editing:** Narayan Gyawali, Melissa Graham, Mohammad Shafiul Alam, Audrey Lenhart, Zhiyong Xi, Gordana Rašić, Nigel W. Beebe, Leon E. Hugo, Gregor J. Devine

**Disclaimer:** The findings and conclusions in this report are those of the authors and do not necessarily represent the official position of the U.S. Centers for Disease Control and Prevention.

